# Simpler and Faster Development of Tumor Phylogeny Pipelines

**DOI:** 10.1101/2021.08.29.458137

**Authors:** Sarwan Ali, Simone Ciccolella, Lorenzo Lucarella, Gianluca Della Vedova, Murray Patterson

**Affiliations:** Department of Computer Science, Georgia State University, Atlanta, GA 30303, USA; Department of Informatics, Systems, and Communications, University of Milano – Bicocca viale Sarca 336, Milano, Italy

**Keywords:** Single Cell Sequencing, Cancer Analysis, Autoencoder, Ridge Regression, Dimensionality Reduction

## Abstract

In the recent years there has been an increasing amount of single-cell sequencing (SCS) studies, producing a considerable number of new datasets. This has particularly affected the field of cancer analysis, where more and more papers are published using this sequencing technique that allows for capturing more detailed information regarding the specific genetic mutations on each individually sampled cell.

As the amount of information increases, it is necessary to have more sophisticated and rapid tools for analyzing the samples. To this goal we developed `plastic`, an easy-to-use and quick to adapt pipeline that integrates three different steps: (1) to simplify the input data; (2) to infer tumor phylogenies; and (3) to compare the phylogenies.

We have created a pipeline submodule for each of those steps, and developed new in-memory data structures that allow for easy and transparent sharing of the information across the tools implementing the above steps.

While we use existing open source tools for those steps, we have extended the tool used for simplifying the input data, incorporating two machine learning procedures — which greatly reduce the running time without affecting the quality of the downstream analysis. Moreover, we have introduced the capability of producing some plots to quickly visualize results.

## 1 Introduction

During the last few years we have witnessed an explosion of computational tools to infer tumor phylogenies (also called cancer progressions) from singlecell sequencing (SCS) data.

Most of the algorithmic research has so far focused on bulk sequencing data for inferring tumor phylogenies, mainly because of the widespread availability and affordability of the next-generation sequencing (NGS) data that are used, producing a large number of tools Strino *et al*. (2013); Jiao *et al*. (2014); Hajirasouliha *et al*. (2014); Yuan *et al*. (2015); Popic *et al*. (2015); Malikic *et al*. (2015); El-Kebir *et al*. (2016); Marass *et al*. (2016); Satas and Raphael (2017); Bonizzoni *et al*. (2018); Toosi *et al*. (2019); Wu (2019). From a computational point of view, the main characteristic of bulk sequencing data is that only the approximate proportion of cells with any given mutation is observable, without distinguishing the cells that carry them. Moreover, each sample contains a mixture of both healthy and cancerous cells — the latter belonging to different and unknown clones — therefore further introducing uncertainty.

More recently, the introduction of single-cell sequencing (SCS) technologies promises to greatly reduce such uncertainty, since the presence or absence of mutation is determined at the level of the cell. Unfortunately, SCS is still much more expensive than bulk sequencing, hence limiting its adoption in practice. Moreover, the quality of the data obtained from SCS is not yet at par with bulk sequencing Kharchenko (2021). In fact, those datasets are affected by some clearly identified problems: (1) doublet cell captures, that is, data originating from two cells instead of one; (2) false negatives from allelic dropout, that is, the presence of a mutation is not detected; and (3) missing values due to low coverage. However, all three of these problems are slowly fading away — the latter two driven by the reduction in cost that allows a higher coverage, and the first due to the development of state-of-the art approaches DePasquale *et al*. (2019) are able to remove such artifacts.

Various methods have been recently developed for inferring tumor phylogenies given current SCS data Jahn *et al*. (2016); Ross and Markowetz (2016); Zafar *et al*. (2017, 2019); Ciccolella *et al*. (2020b); El-Kebir (2018); Singer *et al*. (2018), some of them introducing a hybrid approach of combining both SCS and VAF (variant allele frequency, from bulk sequencing) data Ramazzotti *et al*. (2019); Malikic *et al*. (2017); Salehi *et al*. (2017). To trim the search space and reduce the time needed to infer a phylogeny, most methods rely on the infinite sites assumption (ISA), which essentially states that each mutation is acquired at most once in the phylogeny and is never lost. This assumption leads to a computationally tractable model of evolution called perfect the phylogeny Gusfield (1991); Kimura (1969).

However, some studies Kuipers *et al*. (2017); Brown *et al*. (2017) on cancer data provide strong hints that the ISA does not always hold, the main reason is that cancers usually have large independent deletions on separate branches of the phylogeny Brown *et al*. (2017). When those deletions span a shared locus, we observe multiple deletions of the same mutation.

Relaxing the ISA greatly expands the search space, making it more difficult to develop efficient approaches. For this reason, the number of possible mutation losses is usually bounded in any (more general) model which relaxes the ISA. While the general Dollo model Farris (1977); Rogozin *et al*. (2006) does not impose any restriction on the number of losses, more restricted models are the Dollo*k* and the Dollo-1 (also known as the persistent phylogeny Bonizzoni *et al*. (2012, 2017)).

Even though relaxing the ISA increases the complexity, some methods have appeared, such as TRaIT Ramazzotti et al. (2019), SiFit Zafar *et al*. (2017), SASC Ciccolella *et al*. (2020b) and SPhyR El-Kebir (2018). The latter is extremely relevant to our context, as it introduces the idea of clustering the input SCS matrix to reduce the running time. In Ciccolella *et al*. (2021) this idea is further developed, by devising a clustering method tailored to SCS data — while SPhyR instead relied on the ubiquitous *k*-means McQueen (1967); Anderberg (1973) algorithm. Since SCS data are becoming cheaper to produce, we expect the datasets to increase more rapidly than computing power, as mentioned explicitly in Kharchenko (2021). Therefore including some steps to reduce the instance matrix will become even more common in the next years.

The availability of so many tools for inferring tumor phylogenies, not to mention the parameters that those methods routinely have, means that it is easy to have several different phylogenies from the same SCS dataset, motivating the search for methods that are able to compare and cluster those phylogenies — a large cluster with several highly similar phylogenies is likely to be more reliable, since the underlying evolution is confirmed by multiple methods. In this direction, some methods to measure the distance between two tumor phylogenies have recently been proposed DiNardo *et al*. (2019); Karpov *et al*. (2019); Govek *et al*. (2018); Bernardini *et al*. (2019, 2020); Ciccolella *et al*. (2020a); Jahn *et al*. (2021). While these measures vary greatly in practical applicability, they all express the need for incorporating the evolutionary process into the definition of distance.

Our discussion so far shows that tumor phylogeny inference is crystallizing into different, well-established steps that are combined to obtain a complete tool that starts from an SCS dataset and ends with one or more phylogenies together with some rough idea of their relationships. Still, how to combine those steps is largely ad-hoc, making it more time consuming than is necessary to develop a complete pipeline.

With the goal of making the analysis of cancer data more streamlined, we developed `plastic` (PipeLine Amalgamating Single-cell Tree Inference Components), an integrated and easy to use tool that includes clustering, phylogeny inference, and comparison steps. The `plastic` tool is developed in Python and can be easily integrated into any script or used inside an interactive notebook, such as Jupyter Notebook, to facilitate the reproduction of research results. Currently, `plastic` incorporates the publicly available tools *celluloid, SASC*, and *MP3treesim* — respectively for the clustering, inference, and distance steps — but provides the infrastructure to easily extend it to incorporate any other tool. In fact, `plastic` provides unified in-memory data structures to manage the communication between steps.

Moreover, the current strategies for trimming SCS datasets focus on reducing the number of mutations, while leaving the set of cells unchanged. We have divided the clustering step into two parts: first reducing the number of cells, then reducing the number of mutations. We have explored two strategies for the first part that has been proven to be useful in the past, namely, ridge regression Hoerl and Kennard (1970); Marquardt and Snee (1975) and autoencoder Wang *et al*. (2014).

Ridge regression is a variant of least squares regression, which deals with the trade-off between bias and variance while fitting the regression line. As compared to least squares regression, ridge regression consist of an *l*_2_ penalty on the regression coefficients (see Equation 1). This penalty term in ridge regression helps to introduce some bias, which eventually helps to reduce the variance while fitting the regression line (hence regularizing the fit). In least squares regression, since we do not have the bias, there will be a high variance while fitting the line. Ridge regression has been successfully used in the literature for dimensionality reduction Chen *et al*. (2018); Imakura *et al*. (2019); Liu *et al*. (2019).

Autoencoder is an unsupervised approach based on artificial neural networks. It takes as input the data matrix, encodes it into a latent space (of reduced dimension), and then tries to reconstruct the original input from the latent space. During the reconstruction process, it tries to minimize least squared error. Autoencoder has been successfully used in the literature for dimensionality reduction Ramamurthy *et al*. (2020); Hu and Greene (2018); D’Agostino *et al*. (2018); Abeßer *et al*. (2017). For further detail about ridge regression and autoencoder, please refer to Section 2.1. The idea with both ridge regression and autoencoder is that we can perform downstream clustering (tree inference, etc.) in reduced dimensional data, rather than applying clustering, etc., on the high dimensional data, resulting in reduced downstream runtimes.

We have run two different experiments: one on real data to showcase all features of `plastic`, including its capability to plot clustering of mutations (*i*.*e*., the output of the clustering step) and trees; and the second on simulated data, to assess the reduction in running time stemming from the reduction in the number of cells provided by the two dimensionality reduction strategies mentioned above. The `plastic` tool and all data needed to reproduce the analysis can be found at https://github.com/plastic-phy, including the source code of the Jupyter notebook used for the real data experiment, witnessing the simplicity of our approach. The `plastic` tool is available under the MIT license.

## 2 Methods

We have developed an integrated, modular, and extendable tool for inferring and comparing cancer progressions (also referred to as tumor phylogenies) called `plastic`, which integrates three separate steps into a single program that shares the same data structure among these steps. These three steps are (1) input matrix reduction, (2) tumor phylogeny inference, and (3) tumor phylogeny comparison.

One of the main contributions of our paper is the integration of different tools that usually have specific on-disk input and output file formats, therefore needing a parsing step to process the input, and a dedicated procedure to produce the output. Instead, `plastic` uses in-memory data structures to share all information across the methods.

In particular, `plastic` provides an SCS matrix data structure that is enriched with some additional information that can be shared among the different steps, and a phylogeny tree structure that is used for communication between the phylogeny inference and the phylogeny comparison steps. Such structures are transparent to the user and contribute many additional functionalities without complications from the end-user perspective.

Furthermore, given the nature of `plastic`, we added some graphical capabilities, so that each step of the pipeline can be displayed within an interactive notebook or be exported to separate files.

Finally, notice that each step is optional, therefore allowing the execution of the entire pipeline, or only of a part of it. A schematic of our `plastic` tool is depicted in Figure 1. We now introduce some of the new dimensionality reduction features we have added to `plastic`, as follows.

**Figure 1:**
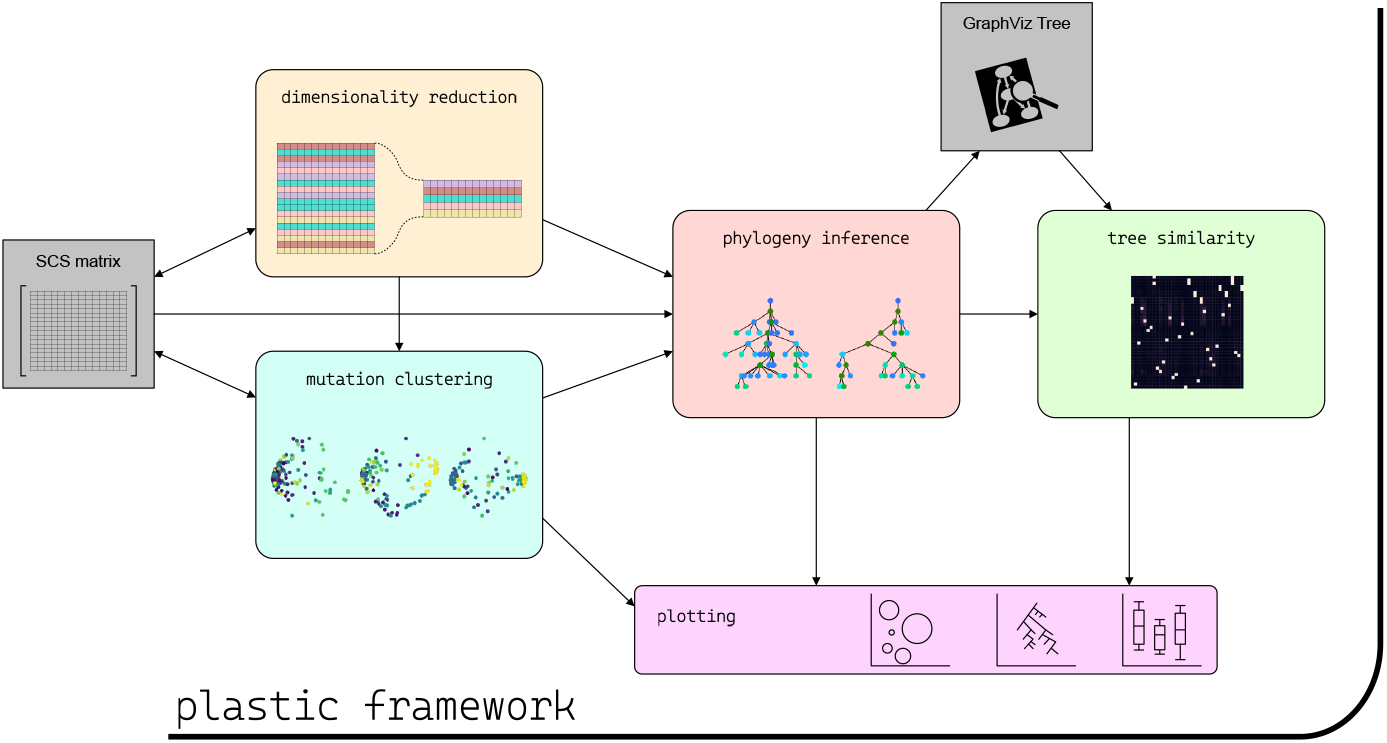
Graphical representation of the `plastic` framework and the interaction between its components and with the input/output files.

### 2.1 Dimensionality Reduction

Dimensionality reduction is a popular approach to enhance the performance of machine learning algorithms and to avoid the problem of the “curse-of-dimensionality” Ali *et al*. (2019a,b). To increase the clustering performance and reduce the runtime for clustering, we use two dimensionality reduction approaches, namely ridge regression Hoerl and Kennard (1970); Marquardt and Snee (1975) and autoencoder Wang *et al*. (2014).

The main goal of ridge regression (RR) is to find a linear function, which models the dependencies between covariate variables and univariate labels. Although ridge regression is an older approach, it is still successfully used in order to reduce the dimensions of current datasets Chen *et al*. (2018); Zhang *et al*. (2010). Ridge regression help to find a subspace, which most compactly expresses the target and rejects other possible but less compact candidates Imakura *et al*. (2019). It works by introducing a bias term — the goal of ridge regression is to increase the bias (by changing the slope of the regression line) in order to improve the variance (generalization capability). The general expression for ridge regression is as follows:

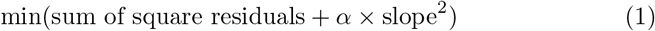

where (*α*×slope^2^) is an *l*_2_ penalty term. Ridge regression gives insights on which independent variables are not informative (independent variables for which we can reduce the slope close to zero). We can eliminate those independent variables to reduce the dimensions of the data. Note that after the dimensionality reduction, we are left with the variables from the actual data and not some latent variables.

Autoencoder (AE) is an unsupervised artificial neural network based approach used for dimensionality reduction Wang *et al*. (2012, 2016). It consists of three the components: encoder; code; and decoder (pictured in Figure 2). The encoder component compresses the input data and produces the code (low dimensional latent-space representation). The decoder component then reconstructs the input only using this low dimensional representation only (hence an unsupervised approach). The low-dimensional data (in the code component) is a compact summary, or compression of the input. This compact data is used as the reduced dimensional representation of the original data. The activation function used for the encoding component is the rectified linear activation function (ReLU), while we used the sigmoid function for the decoding. The loss function that we used in our experiments is the least squared error. The optimizer that we use is *Adadelta* Polic *et al*. (2019), a stochastic gradient descent method which is based on adaptive learning rate per dimension. It is a popular optimizer used in autoencoder because it avoids the continual decay of learning rates throughout training. It also helps to decide the global learning rate, hence not requiring to select it manually.

**Figure 2:**
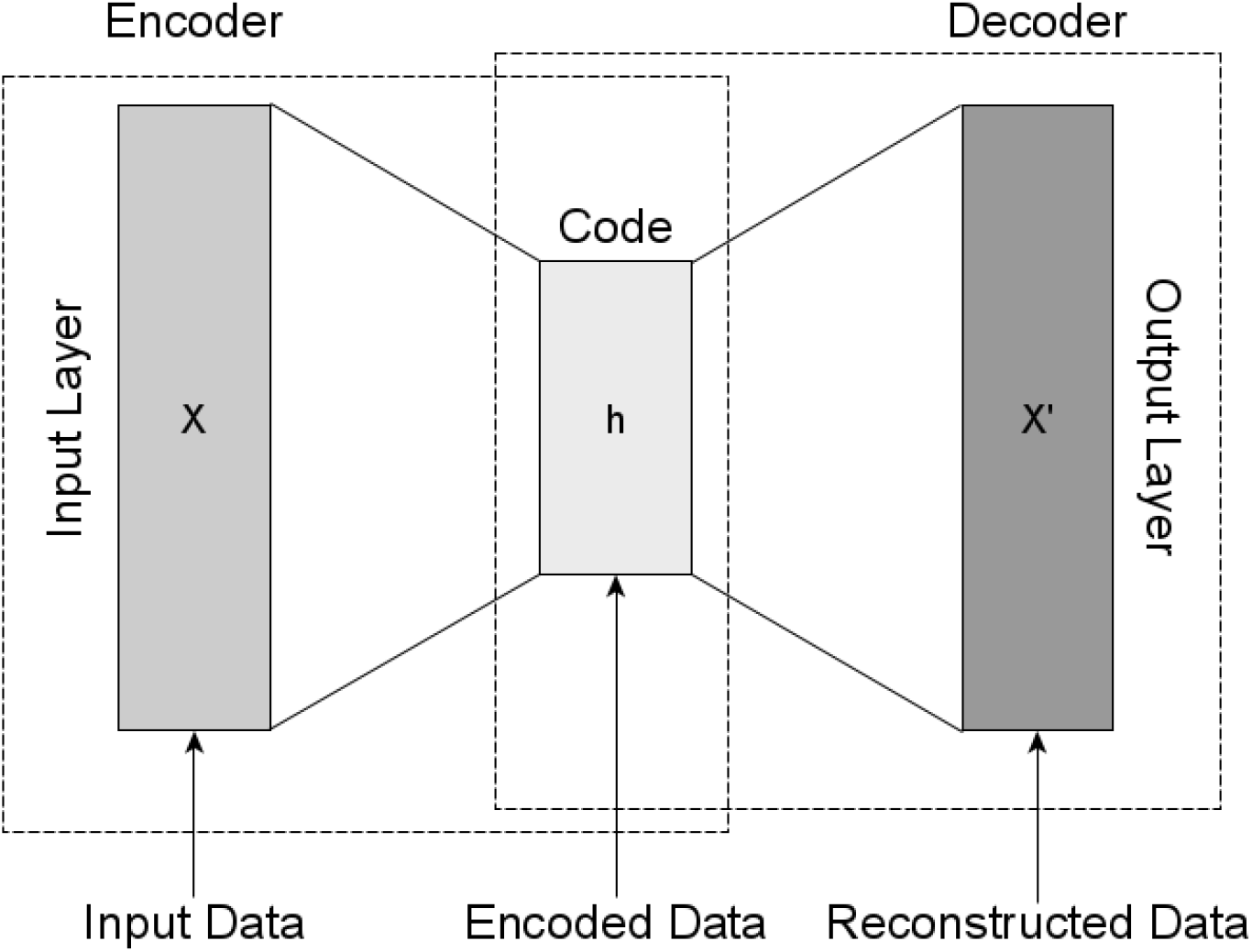
The autoencoder architecture. The encoder takes input data and maps it to a latent space (code component) of reduced dimension. The decoders code back the latent representation to the output. This architecture learns to compress data by reducing the reconstruction error (least squared error).

## 3 Results

Here we present our results on both real and simulated data.

### 3.1 Real data

As a result, we show a complete workflow example on a real medulloblastoma dataset Hovestadt *et al*. (2019). In particular, we will explore a full pipeline analysis where multiple samples are taken into account. For each sample: mutations are clustered using *celluloid*; phylogeny trees are reconstructed using *SASC*; and finally, the patients are clustered using *MP3treesim* as similarity measure between them.

The sequencing study consists of 36 patients that we want to cluster into different subtypes. To simulate a real case scenario, where such information is not available, we started from SCS binary matrices, one for each patient, and we clustered mutations using *celluloid* (*k* = 50), for which we can see a summary in Figure 3.

**Figure 3:**
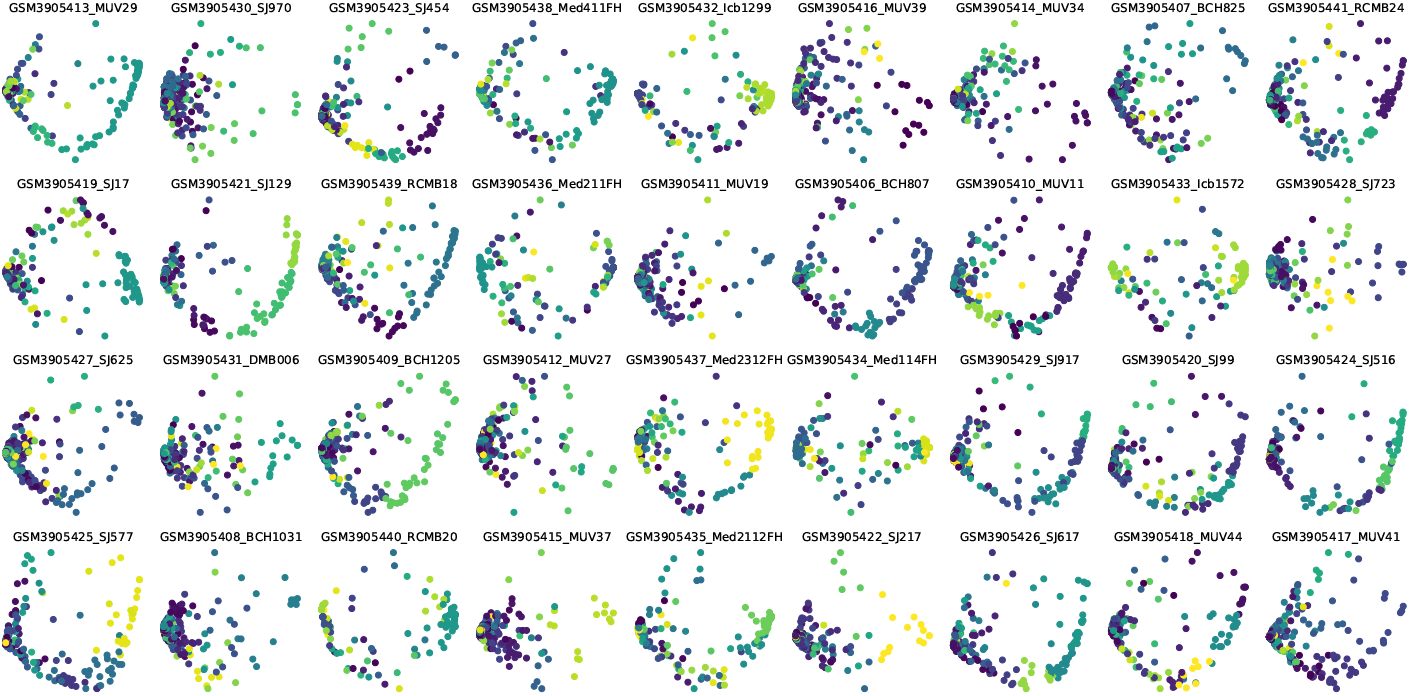
Mutations clustered on SCS data computed by the *celluloid* submodule and displayed by `plastic` for the 36 medulloblastoma patients in the dataset of Hovestadt *et al*. (2019).

After the first preprocessing step, we utilized the *SASC* submodule of `plastic` to infer the evolutionary trees of each patient. As a proof of concept we run it using the following parameters *α* = 0.25, *β* = 1 *×* 10^*−*4^, *k* = 0, *d* = 0 for whose definitions we refer to *SASC* ‘s manuscript Ciccolella *et al*. (2020b). Once computed we used the newly-added plotting feature to display the phylogenies using the *SASC-viz* plotting feature to prettify the trees; the result is shown in Figure 4.

**Figure 4:**
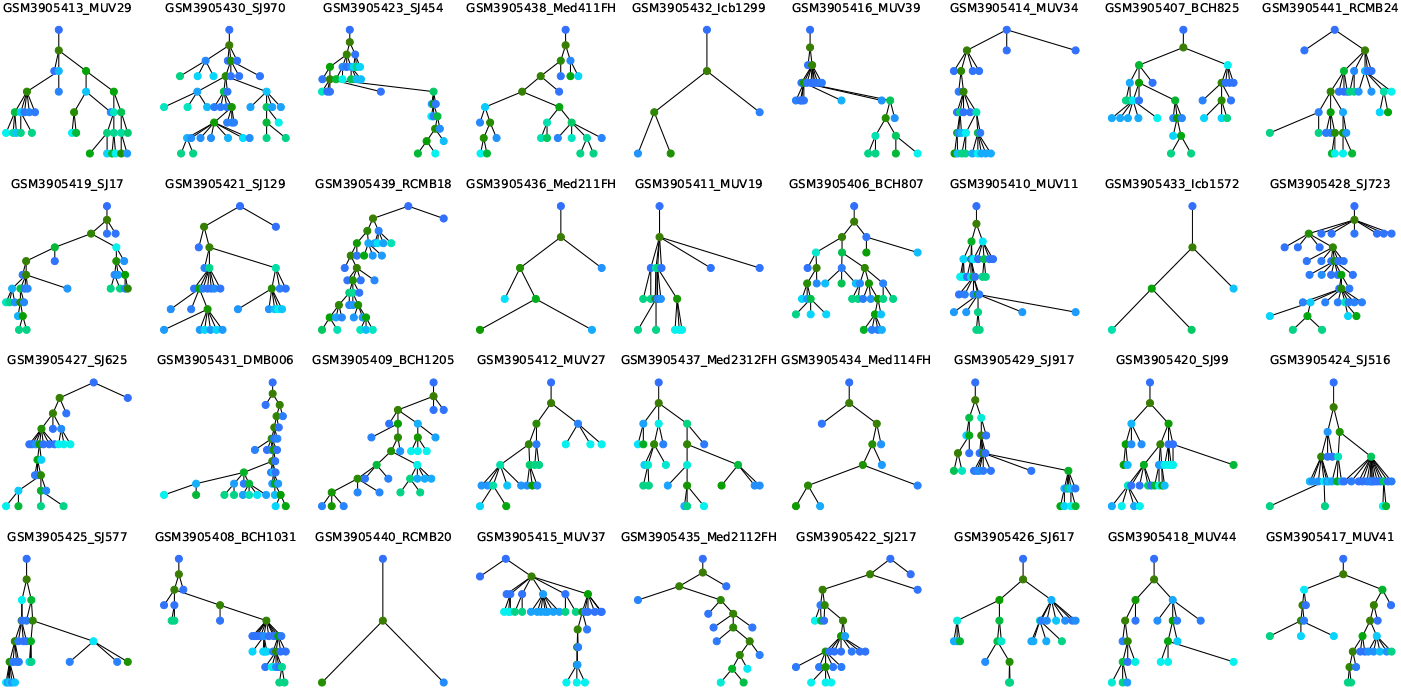
Trees computed by the *SASC* submodule and displayed by `plastic` for the 36 medulloblastoma patients in the dataset of Hovestadt *et al*. (2019).

As the last step, we then used the *MP3treesim* submodule to compute a matrix of similarity-scores between all the trees and then used it to cluster the patients according to a hierarchical clustering method. We then displayed the final clustering of the patients in Figure 5.

**Figure 5:**
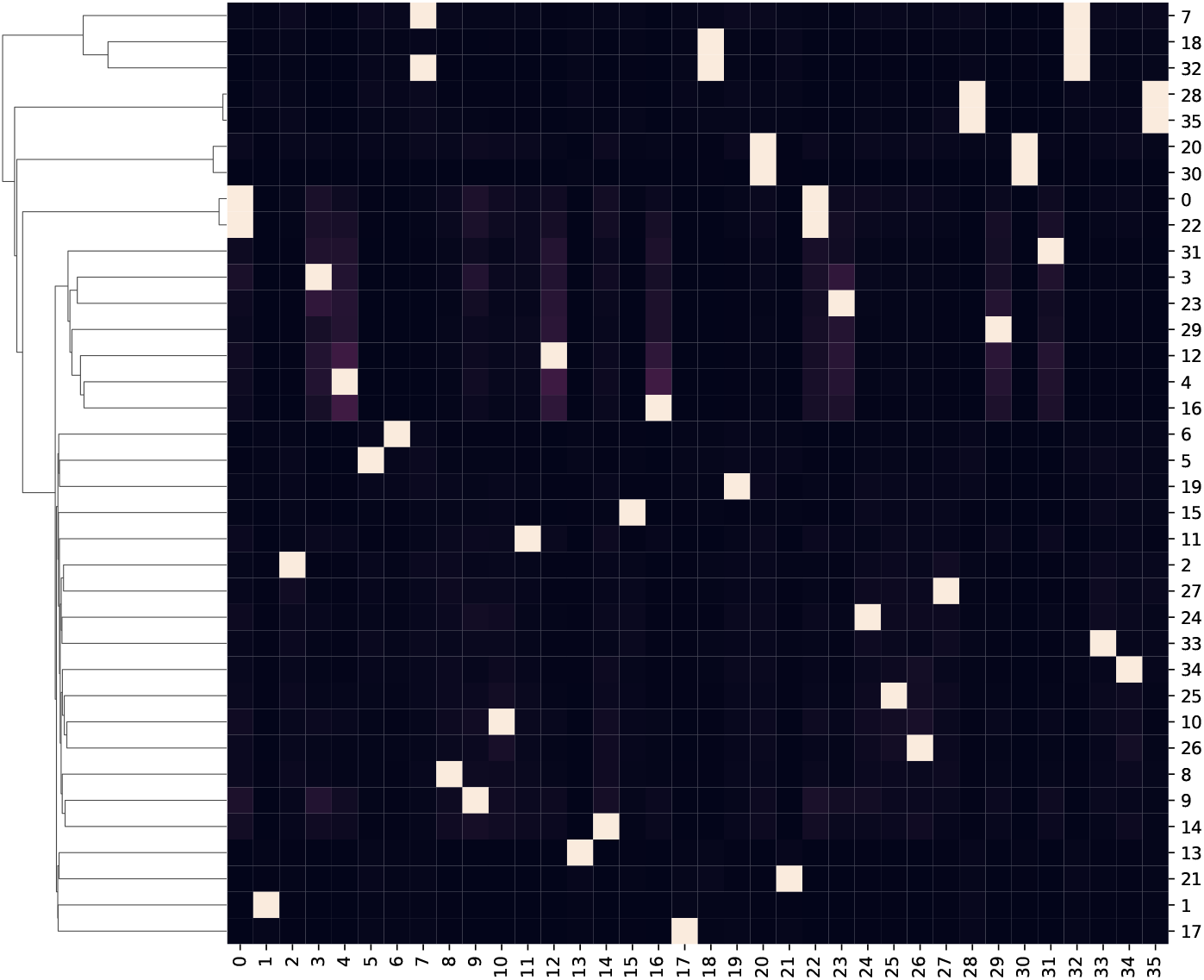
Clustermap based on the similarity between the trees computed by the mp3 submodule of `plastic` for the 36 medulloblastoma patients in the dataset of Hovestadt *et al*. (2019).

### 3.2 Simulated data

To evaluate the performance of the input matrix reduction methods that we present here, we designed a two-fold experiment on synthetic datasets. We first measure the quality of the matrix obtained from the reduction step — in this case we have a ground truth that we can use for the evaluation. Second, we evaluate the accuracy and runtime of the downstream phylogeny inference tools when given as input the reduced instance from the first step, to assess that our reduction step is actually useful. This is similar to the experimental approach taken in Ciccolella *et al*. (2021).

The simulated data are generated as follows. First we generate a random tree topology on *s* nodes, each representing a tumor clone, by first creating a root (the germline) and then iteratively attaching the *s−*1 remaining nodes uniformly at random to any other node in the tree. The nodes are then randomly labeled with *m* mutations — each mutation understood as being acquired at the node that it labels. Then, a total of *n* cells are randomly associated to the *s* nodes. Each cell harbours all of the mutations on the path in the tree from the root to the node that the cell labels. A binary *n × m* matrix *M* is then obtained from the cells, where *M* [*i, j*] = 1 if cell *i* harbours mutation *j*, otherwise *M* [*i, j*] = 0. Noise is then added to this matrix according to the false negative, false positive and missing value rates, to simulate a real single cell sequencing experiment. Each of the *s* nodes is therefore considered as a natural (true) cluster of the simulated dataset.

We design three experiments, each with 100, 200 and 300 cells, respectively. In each experiment, we generate 50 simulated datasets with its corresponding number of cells, using the procedure mentioned above. In all three experiments, the number *s* of clones is 20, and the number *m* of mutations is 1000.

To each dataset of our experiments, we first performed dimensionality reduction in the cells using both the ridge regression and autoencoder techniques mentioned in Section 2. The number of dimensions selected in case of autoencoder are 50, 100, and 150 for experiments 1, 2 and 3, respectively. This number of dimensions, in each case, is selected empirically when the least squared error for the loss function is minimized. The loss function gives us the error value when autoencoder tries to reconstruct the input in the decoder component. The main goal in this case is to reconstruct the input as accurately as possible, giving rise to the respective numbers of dimensions above. The number of dimensions selected in the case of ridge regression is different for each dataset because ridge regression is a data driven technique, however the average number of dimensions for each of experiments 1, 2 and 3 is roughly 49, 97, and 146, respectively (see Table 1). Following dimensionality reduction in the cells, we then clustered the mutations using *celluloid*.

**Table 1:** Number of cells selected by ridge regression from the experiments^*a*^

| Experiment | mean ± SD |
| --- | --- |
| 1 | 48.7 ± 2.51 |
| 2 | 97.0 ± 4.71 |
| 3 | 146 ± 5.85 |

<sup>a</sup> The mean ( $\mu$ ) ± standard deviation ( $\sigma$ ) of the number of cells selected from the 50 simulated datasets after applying dimensionality reduction using ridge regression. Calculations reported to three significant digits. The original (full) table, from which these aggregates were performed, can be found at https://github.com/plastic-phy/plastic/blob/master/data/cells-ridge.csv

#### 3.2.1 Evaluating input matrix reduction

Because we are performing dimensionality reduction only on the cells, we can still use the same measures of precision and recall used in Ciccolella *et al*. (2021), which are based on how mutations are clustered together, as follows:

##### Precision

measures how well mutations are clustered together. For each pair of mutations appearing in the same clone in the simulated tree, we check if they are in the same cluster, resulting in a *true positive* (*TP*). For each pair of mutations clustered together that are not in the same clone, we encounter a *false positive* (*FP*). The value of the precision is then calculated with the standard formula: 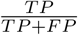

##### Recall

measures how well mutations are separated. For each pair of mutations in the same clone, we now also check if they are not in the same cluster, resulting in a *false negative* (*FN*). The recall is then calculated as: 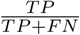

Notice that, just as in Ciccolella *et al*. (2021), we are mostly interested in obtaining a high precision. The reason for focusing on obtaining high precision is that since cancer phylogeny inference algorithms can later cluster together mutations — for example, by assigning them to the same node or the same non-branching path — however they cannot separate mutations that have been erroneously clustered together. It is for this same reason that the number (*k*) of clusters is carefully chosen with high precision in mind.

On the other hand, ridge regression does not allow to determine a priori the number of cells that will be obtained. For this reason we report, in Table 1, the distributions^1^ of the actual number of cells obtained.

Figures 6, 7, and 8 correspond to the precision and recall of the clusterings of the mutations obtained after first applying dimensionality reduction in the cells of experiments 1, 2 and 3, respectively.

**Figure 6:**
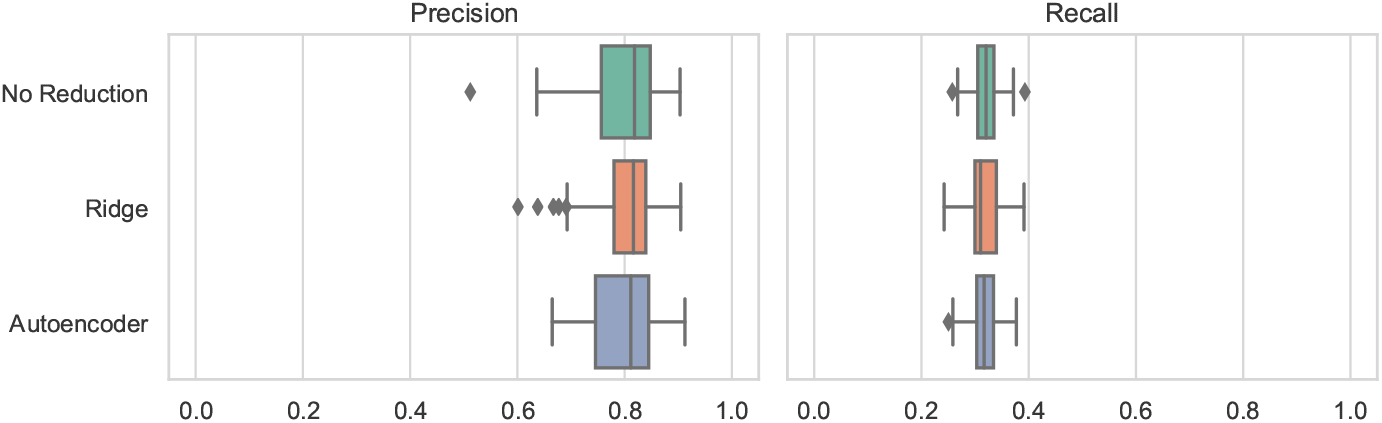
Precision and recall results of the clusterings of the mutations obtained for experiment 1, generated with 1000 mutations and 100 cells. The first row corresponds the results obtained by clustering the mutations to *k* = 100 clusters using *celluloid* (no dimensionality reduction). The second and third rows correspond to first applying a dimensionality reduction step on the 100 cells, using ridge regression and autoencoder, respectively, followed by the clustering of the 1000 mutations to *k* = 100 clusters (in the reduced number of cells) using *celluloid*.

**Figure 7:**
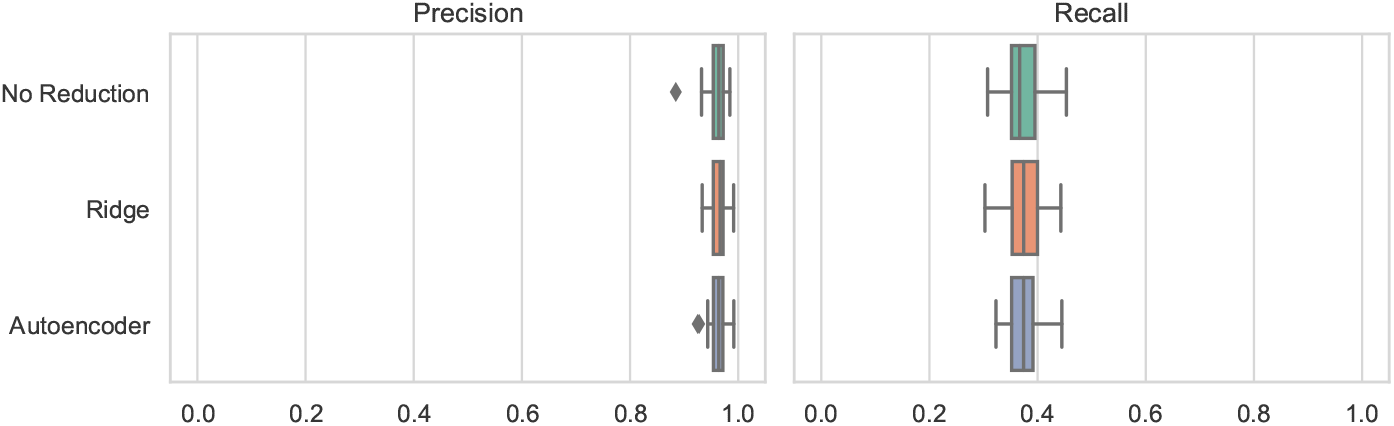
Precision and recall results of the clusterings of the mutations obtained for experiment 2, generated with 1000 mutations and 200 cells. The first row corresponds the results obtained by clustering the mutations to *k* = 100 clusters using *celluloid* (no dimensionality reduction). The second and third rows correspond to first applying a dimensionality reduction step on the 200 cells, using ridge regression and autoencoder, respectively, followed by the clustering of the 1000 mutations to *k* = 100 clusters (in the reduced number of cells) using *celluloid*.

**Figure 8:**
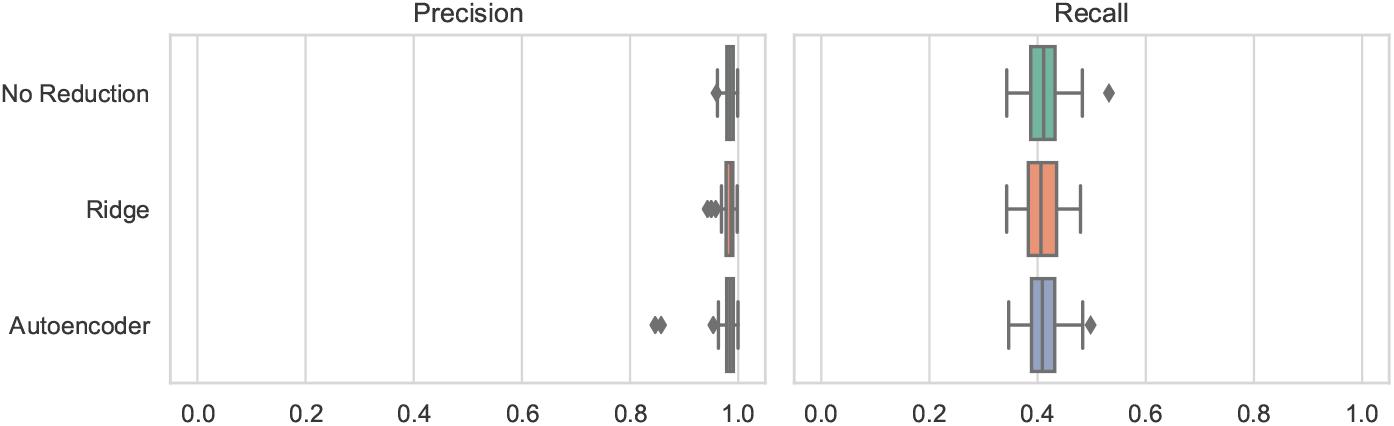
Precision and recall results of the clusterings of the mutations obtained for experiment 3, generated with 1000 mutations and 300 cells. The first row corresponds the results obtained by clustering the mutations to *k* = 100 clusters using *celluloid* (no dimensionality reduction). The second and third rows correspond to first applying a dimensionality reduction step on the 300 cells, using ridge regression and autoencoder, respectively, followed by the clustering of the 1000 mutations to *k* = 100 clusters (in the reduced number of cells) using *celluloid*.

#### 3.2.2 Evaluating the effect of the reduction step on downstream phylogeny inference

The goal of dimensionality reduction (in the cells) and clustering (of the mutations) is to allow for a significant decrease in the runtime of the phylogeny inference, the most expensive step in the pipeline — as long as it does not worsen the accuracy. The idea is that, since the reduced matrix has much fewer rows and columns, using it in place of the original matrix as input to the phylogeny inference should result in much lower runtimes, at the risk of decreasing the accuracy of downstream analysis, due to errors introduced in the dimensionality reduction and clustering steps. The main observation is that, if the clusters are highly precise, the decrease of the accuracy of the downstream inference is negligible or small. Assessing this fact is the main goal of our experimental analysis, based on the measures used in Ciccolella *et al*. (2020c,b):

##### Ancestor-descendant accuracy

For each pair (*m*_1_, *m*_2_) of mutations such that *m*_1_ is an ancestor of *m*_2_ in the ground truth, we check whether *m*_1_ is an ancestor of *m*_2_ also in the inferred tree (*TP*) or whether it is not (*FN*). Moreover, each pair (*m*_1_, *m*_2_) of mutations such that *m*_1_ is an ancestor of *m*_2_ in the inferred tree but not in the ground truth, we consider that pair a false positive.

##### Different lineages accuracy

Similarly to the previous measure, we check whether mutations in different branches (*i*.*e*., neither is an ancestor of the other) are correctly inferred or if any pair of mutation is erroneously inferred in different branches.

Since these new dimensionality reduction techniques are built into the `plastic` framework, one can customize a cancer phylogeny inference task by:

1. choosing any (or none of the) dimensionality reduction steps from ridge regression or autoencoder; followed by
2. clustering (or not) with *celluloid*; and finally
3. phylogeny inference with *SASC* in either the reduced set of cells, or not.

An extensive study of how clustering can reduce the runtime (and sometimes improve accuracy) of the downstream phylogeny inference appears in Ciccolella *et al*. (2021), and so here we explore the further gains in runtime and/or accuracy that might be achieved by adding a dimensionality reduction step. Hence, in our experiments, we try three different choices for the reduction step — ridge regression, autoencoder, and no dimensionality reduction to produce a matrix to be given to *celluloid* (notice that *SASC* will receive the matrix on the original set of cells, but with the clustered mutations). Moreover, when applying ridge regression, we also provide the reduced set of cells to *SASC*, hence resulting in four different settings — when no dimensionality reduction is performed, the reduced and the original sets of cells are the same, while autoencoder produces reduced matrices in its learned latent tensor space, which is not an SCS matrix that can be fed to *SASC*. We always perform the clustering step, because 1000 mutations is prohibitive for the downstream phylogeny inference (see Ciccolella *et al*. (2021)).

Figures 9, 10 and 11 report the ancestor-descendant and different lineages accuracy measures of the trees obtained by running `plastic` with the four settings mentioned above. From the results, it is clear that when dimensionality reduction is followed by inference in the original set of cells, there is no noticeable loss in accuracy. In these two cases, the clustering is still performed in the reduced set of cells, providing a speedup at no cost. On the other hand, we see a slight loss of accuracy in the ancestor-descendant measure when ridge regression is applied, and the resulting reduced set of cells is used also in the inference step.

**Figure 9:**
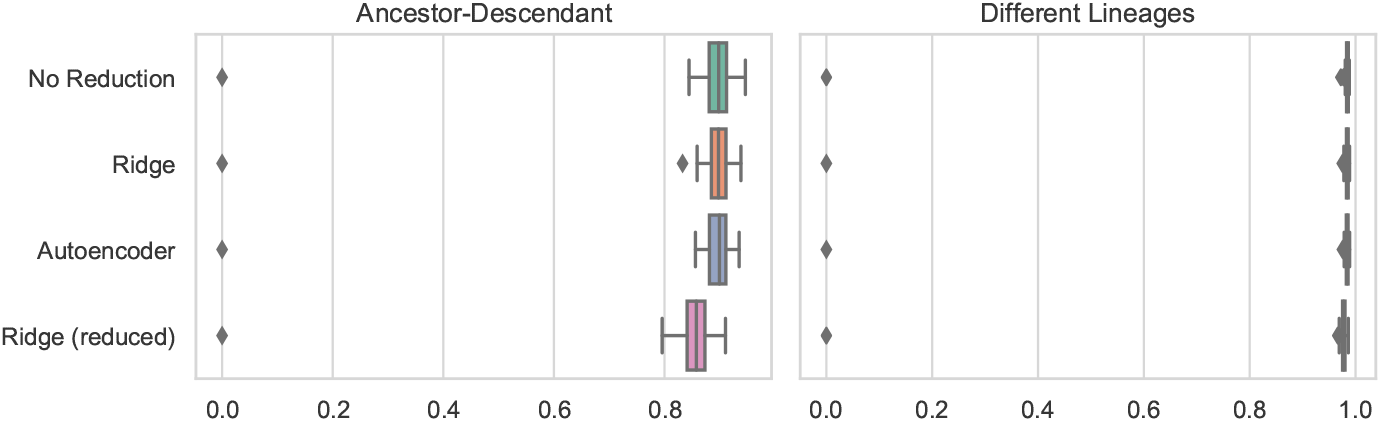
Ancestor-descendant and different lineages accuracies of the trees obtained by running `plastic` on the data of experiment 1. From the top, the four rows correspond to the settings: no dimensionality reduction, just clustering — No Reduction —; ridge regression + clustering in the original set of cells — Ridge —; autoencoder + clustering in the original set — Autoencoder —; and ridge regression + clustering in the reduced set — Ridge(reduced).

**Figure 10:**
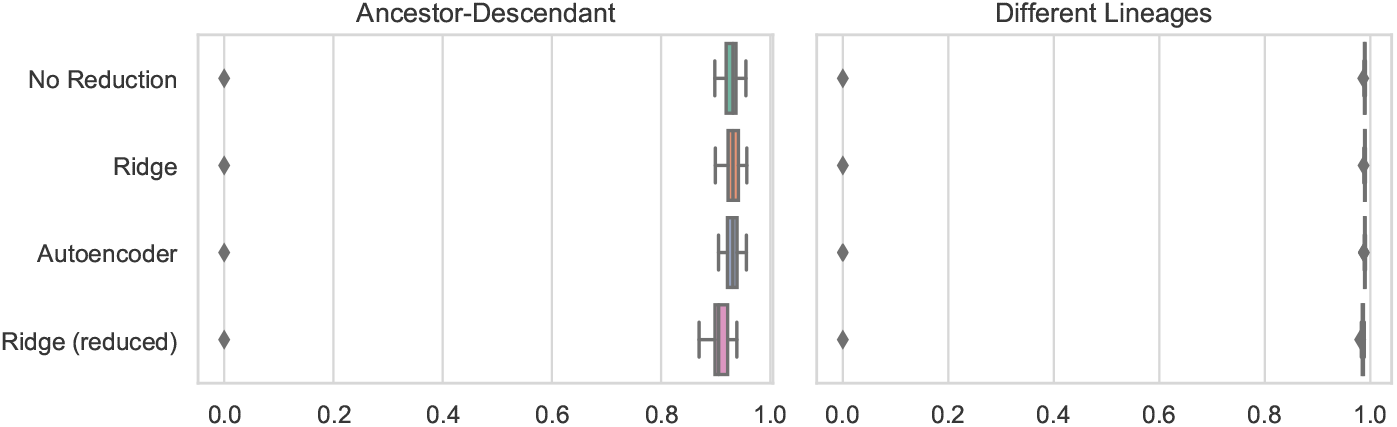
Ancestor-descendant and different lineages accuracies of the trees obtained by running `plastic` on the data of experiment 2. From the top, the four rows correspond to the settings: no dimensionality reduction, just clustering — No Reduction —; ridge regression + clustering in the original set of cells — Ridge —; autoencoder + clustering in the original set — Autoencoder —; and ridge regression + clustering in the reduced set — Ridge(reduced).

**Figure 11:**
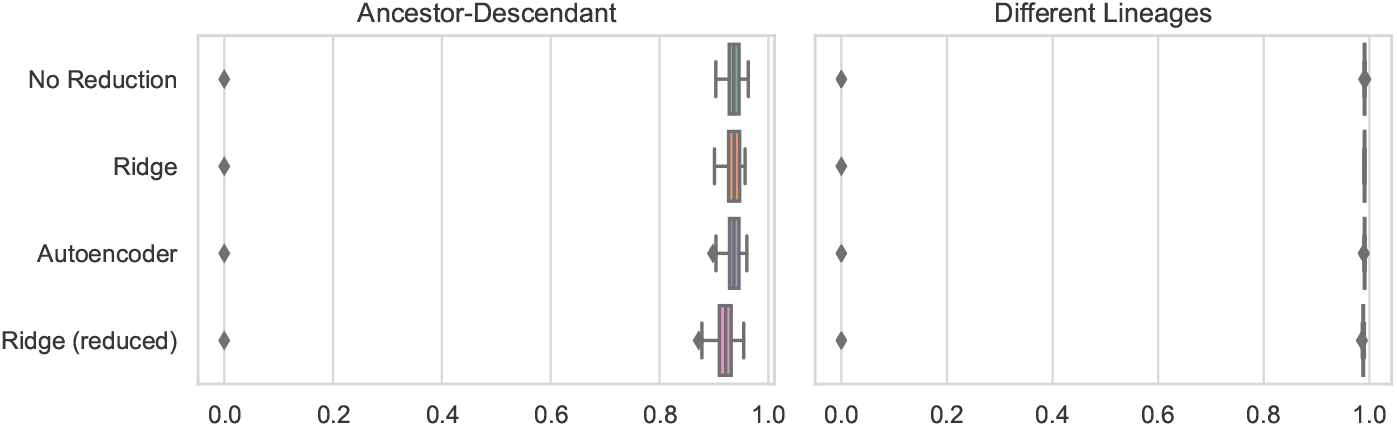
Ancestor-descendant and different lineages accuracies of the trees obtained by running `plastic` on the data of experiment 3. From the top, the four rows correspond to the settings: no dimensionality reduction, just clustering — No Reduction —; ridge regression + clustering in the original set of cells — Ridge —; autoencoder + clustering in the original set — Autoencoder —; and ridge regression + clustering in the reduced set — Ridge(reduced).

We have investigated the speedup provided by ridge regression. In Figure 12 are represented the running times of *celluloid* with and without first applying ridge regression, as a function of the number of mutations. More precisely, we take the first columns of the datasets of Experiment 1 (the number of mutations is the *x* axis).

**Figure 12:**
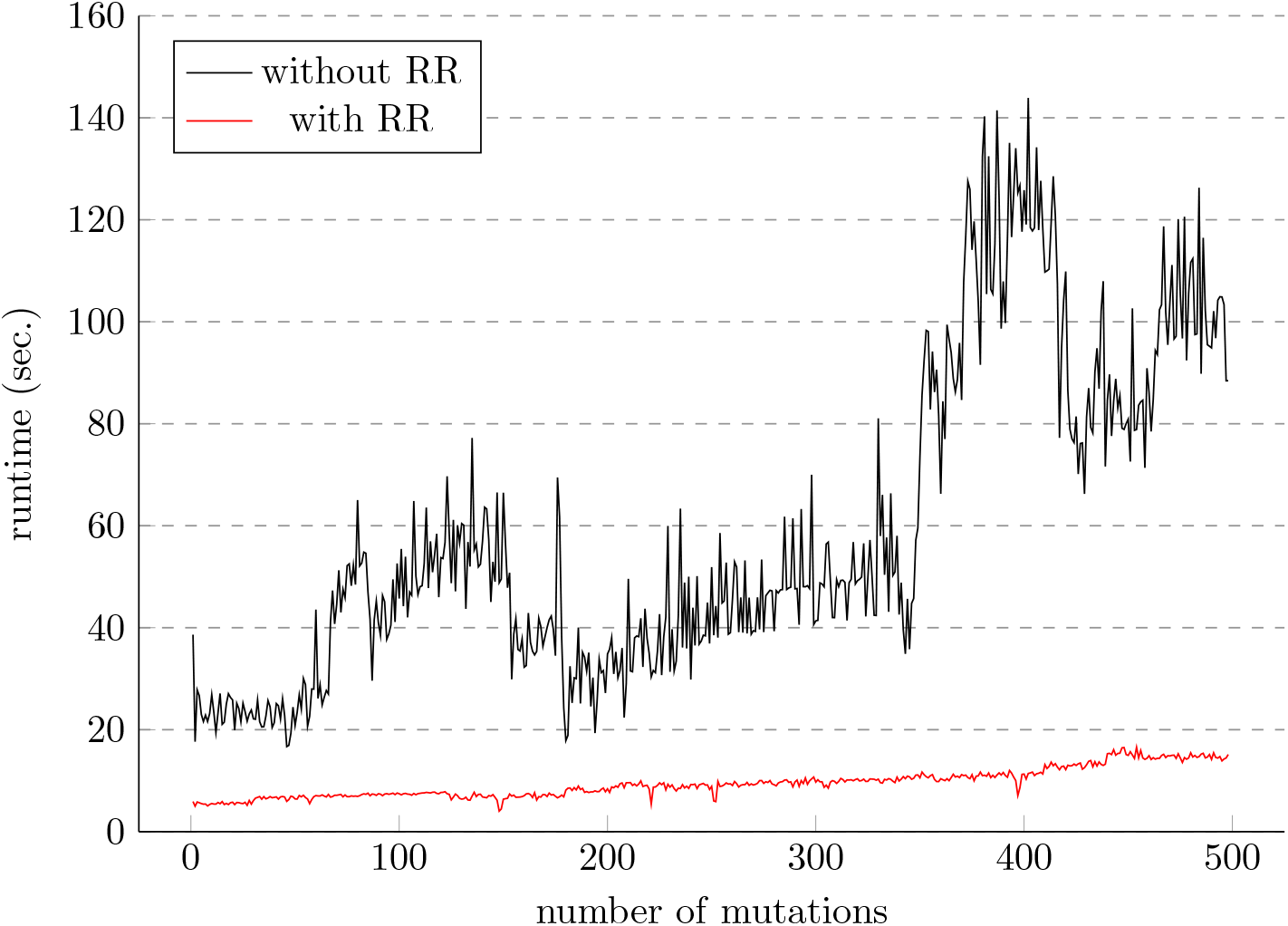
Runtime analysis, as a function of the number of mutations, of the clustering step (*celluloid*) without dimensionality reduction using ridge regression (in black), and with ridge regression (in red).

Finally, in Table 2 we report the running times of *SASC* when ridge regression is applied, and the resulting reduced set of cells is used also in the inference step, that is, the case where we have observed a slight loss of accuracy in the ancestor-descendant measure. In this case the phylogeny inference is up to 100 times faster, making therefore possible to run *SASC* on more mutations.

**Table 2:** Runtimes of *SASC* on the experiments^*a*^

| Experiment | **celluloid** | **RR + celluloid** |
| --- | --- | --- |
| 1 | $37.7 \pm 24.3$ | $0.414 \pm 0.0710$ |
| 2 | $44.3 \pm 21.2$ | $0.832 \pm 0.0682$ |
| 3 | $34.2 \pm 25.6$ | $1.31 \pm 0.112$ |

<sup>a</sup> The mean ( $\mu$ ) ± standard deviation ( $\sigma$ ) of the runtime of *SASC* on the 50 datasets of each of the three experiments (on the original set of cells), after having either: clustered the mutations of the input matrix using *celluloid* (**celluloid**); or performed dimensionality reduction on the cells using ridge regression, followed by clustering the mutations with *celluloid* (**RR + celluloid**). Times are reported in hours, and to three significant digits.

## 4 Discussion

Given the large amount of SCS cancer studies that are being published, there is a need for a fast and easy-to-use framework that allows to perform the needed analyses. We developed `plastic` with this goal in mind, a pipeline composed of different submodules that allow for dimensionality reduction on SCS cells, clustering of SCS mutations, inference of tumor phylogenies, comparison of such trees, and convenient plotting of the results.

Each of the submodules can be used independently or they can be used in conjunction with each other to create complex operations, due to special data structures developed for the interaction between different methods.

The `plastic` pipeline is open-source and available at https://github.com/plastic-phy along with extensive documentation and a Jupyter Notebook that replicates the real-case scenario depicted in Section 3.1.

Future improvements for `plastic` would be to include more tools into the pipeline to extend the breadth of analyses available and to provide additional algorithmic alternatives for the same task. Moreover, we could incorporate more steps, such as the removal of doublets, to streamline the entire pipeline even more.

## Acknowledgments

We thank Paola Bonizzoni and Iman Hajirasouliha for many useful discussions on the topic.

## Footnotes

1 note that the variance of all three distributions is small

